# Phosphorus Bioaccessibility of Processed Soy and Pulse Protein Products Measured by *In Vitro* Simulation of Human Digestion

**DOI:** 10.1101/2025.05.13.653545

**Authors:** Kendal M. Burstad, Alexandria Fons, Abigail R. Kisch, Dennis P. Cladis, Kathleen M. Hill Gallant

## Abstract

**Objective:** Plant protein is gaining interest for dietary phosphorus management in CKD due to its potential benefits, including possible lower phosphorus bioaccessibility. However, data on phosphorus remain limited, particularly for processed plant protein products. This pilot study aimed to compare phosphorus bioaccessibility of processed plant protein products with their animal protein counterparts, using *in vitro* simulation of adult human digestion.

**Methods:** Soy protein (n=16), pulse protein (n=17), and animal protein (n=13) products representing food categories of beef, dairy, sausage/bacon, chicken/turkey were evaluated for phosphorus bioaccessibility using *in vitro* digestion based the INFOGEST protocol, followed by dialysis of the final digesta. Pre-digestion samples, final digesta, and final dialysate were analyzed for phosphorus content to calculate total and bioaccessible phosphorus and percent phosphorus bioaccessibility.

**Results:** Average percent phosphorus bioaccessibility of all processed soy and pulse products studied ranged from 32-100%, while animal products ranged from 81-100%. Average bioaccessible phosphorus and bioaccessible phosphorus-to-protein ratio were lower for many soy and pulse protein beef and chicken/turkey alternatives, soy protein milk alternatives, and pulse protein sausage alternatives than their animal protein counter products.

**Conclusion:** Some soy and pulse protein products offered lower bioaccessible phosphorus per 100g serving and per gram of protein than their animal protein counterparts. Thus, this pilot study suggests that certain processed plant protein products may be acceptable to include in a diet for phosphorus management in CKD. However, more data on phosphorus bioaccessibility in a larger number and variety of processed plant-based protein products is needed to better inform individualized guidance.

## Introduction

Plant protein for dietary management of chronic kidney disease (CKD) is of growing interest due to several proposed benefits,^1,2^ including the potential for lower phosphorus bioavailability.^3^ Based on limited data or opinion only, both the Kidney Disease Outcomes Quality Initiative (KDOQI) Nutrition in CKD 2020 Update^4^ and Kidney Disease Improving Global Outcomes (KDIGO) 2017 CKD-Mineral and Bone Disorder (CKD-MBD) Update^5^ suggest considering phosphorus source (animal, plant, or additive) when making dietary recommendations for phosphorus management for people with CKD. These are based on evidence that suggests phosphorus bioavailability is lowest from plant sources, followed by animal sources, and highest from inorganic phosphate (P) additives.^6^ Approximately 50-82% of total phosphorus in plants is in the form of phytic acid,^7,8^ which has limited accessibility for absorption in humans,^9,10^ and thus may reduce overall phosphorus burden.

Although often used interchangeably, ‘bioavailability’ and ‘bioaccessibility’ are distinct concepts (**Box 1**). The commonly cited estimates of phosphorus ‘bioavailability’ (e.g., plant sources: ∼10-50%,^3,6^ animal sources: ∼40-60%,^6^ inorganic P additives: up to ∼100%),^6^ are more accurately estimates of phosphorus ‘bioaccessibility’, the amount of phosphorus digested and in a form that *could* be absorbed.^11,12^ Phosphorus bioaccessibility is a major component and limiting determinant of bioavailability;^11^ but phosphorus bioavailability also depends on the absorption capacity of the intestine.^13^ Because absorbed phosphorus is not known to require further metabolism prior to its utilization by body tissues, we can reasonably consider absorbed phosphorus as bioavailable. However, phosphorus bioaccessibility should not be confused for phosphorus absorption.

Importantly, measuring phosphorus bioavailability in humans is challenging, involving controlled-feeding metabolic balance studies and/or absorption studies using phosphorus isotope tracers.^13,14^ Urine phosphorus is a frequently used biomarker of phosphorus absorption, but urine phosphorus is also affected by renal phosphate clearance and bone turnover rates, which are particularly confounding in people with CKD. To our knowledge, there are no metabolic balance or absorption studies in humans that have investigated phosphorus bioavailability comparing different dietary sources. Contrarily, phosphorus bioaccessibility can be measured more readily by *in vitro* simulation of human digestion. While care should be taken to avoid misinterpreting bioaccessibility as phosphorus absorption (e.g., 100% bioaccessibility ≠ 100% absorption), bioaccessibility studies can provide valuable information on the amount of phosphorus released from different food matrices, which can help guide food choices for dietary phosphorus management.

The limited number of studies on phosphorus bioaccessibility of foods generally substantiate the concept that percent phosphorus bioaccessibility (%PB) increases from plant to animal to inorganic sources, but within groups of foods, there are wide and overlapping ranges: ∼6-42% for legume/seed sources,^15^ ∼29-99% for grain sources,^15,16^ ∼70-107% for meat sources,^17^ ∼49-111% for dairy sources,^17^ and ∼84-100% from inorganic sources.^15^ Additionally, these studies^15–17^ were conducted over a decade ago, prior to the emergence of many current processed plant protein products and were mostly limited to raw/uncooked products. Importantly, processing and cooking methods can affect overall phosphorus content and bioaccessibility of animal and plant protein foods.^18–21^

In addition to the KDOQI^4^ and KDIGO^5^ statements on sources of dietary phosphorus, the KDOQI 2020 nutrition guidelines^4^ provide additional commentary to “advise choosing natural foods that are lower in bioavailable phosphorus” and “advise choosing commercial food items prepared without phosphorus-containing food additives.” This is based on the premise that highly processed (ultra-processed) foods versus “natural” (unprocessed/minimally processed) foods would be more likely to include inorganic P additives with higher bioaccessibility. These foods are also more likely to have other undesirable nutritional attributes such as higher sodium and fat, added sugar, or lower fiber. Certainly, without the necessary data for more precise and nuanced guidance, this commonsense approach is prudent. However, processed foods can also confer benefits that may include convenience, shelf-stability, texture, flavor, and other sensory attributes that have potential to improve nutrition, independence, and quality of life for individuals with diverse needs.^22,23^ However, phosphorus bioaccessibility data for currently available processed plant protein products is absent from the literature. Therefore, the aim of this pilot study was to determine phosphorus bioaccessibility of processed plant protein products (soy and pulse) in comparison to animal protein counterparts using *in vitro* simulated human digestion methods.

## Methods

Further method details are provided in the **Supplementary File**. Briefly, 46 products including 17 pulse protein, 16 soy protein and 13 animal protein products from the categories of beef, dairy (milk, cheese, and yogurt), chicken and turkey, and sausage and bacon were selected for evaluation (**Supplementary Table 1**). Products were prepared according to package directions, frozen, freeze-dried, and stored until analysis (**Supplementary Table 2**). Freeze-dried food samples (pre-digestion) were dry-ashed and analyzed for total phosphorus content by microwave plasma atomic emission spectroscopy (MP-AES) (Agilent Technologies, model 4210) (**Supplementary Methods**). Additional minerals (calcium, magnesium, potassium, and sodium) were also measured, and data are provided in **Supplementary Table 4**.

Freeze-dried food products were digested by *in vitro* simulation of human digestion according to the standardized INFOGEST protocol^24^ with modifications to determine phosphorus bioaccessibility (**Figure 1** and **Supplementary File**). Briefly, the *in vitro* digestion protocol consisted of a 2-minute oral phase, 2-hour gastric, and 2-hour intestinal phase all at 37°C with adjustments in each phase for appropriate pH, digestive fluids and enzymes. Following *in vitro* digestion, 2-3g of final digesta from each sample were dry-ashed and analyzed for phosphorus content by MP-AES and 5 mL of final digesta were dialyzed against ultrapure water using a 500-1000 Dalton molecular weight cut off dialysis membrane device (float-a-lyzer® G2, Repligen G235051). After dialyzing for 30-hours, dialysates were analyzed for phosphorus content by MP-AES. Phosphorus bioaccessibility was calculated as the % dialyzable phosphorus in the digesta for each sample. Any %PB value greater than 100 was censored to the highest possible value of 100%. Bioaccessible phosphorus (BioP, mg) was calculated from the %PB multiplied by total phosphorus content. Data are reported on an “as prepared” basis for all food products. The bioaccessible and total phosphorus content of each product is reported as mg per 100g, unless otherwise indicated.

**Figure 1:**
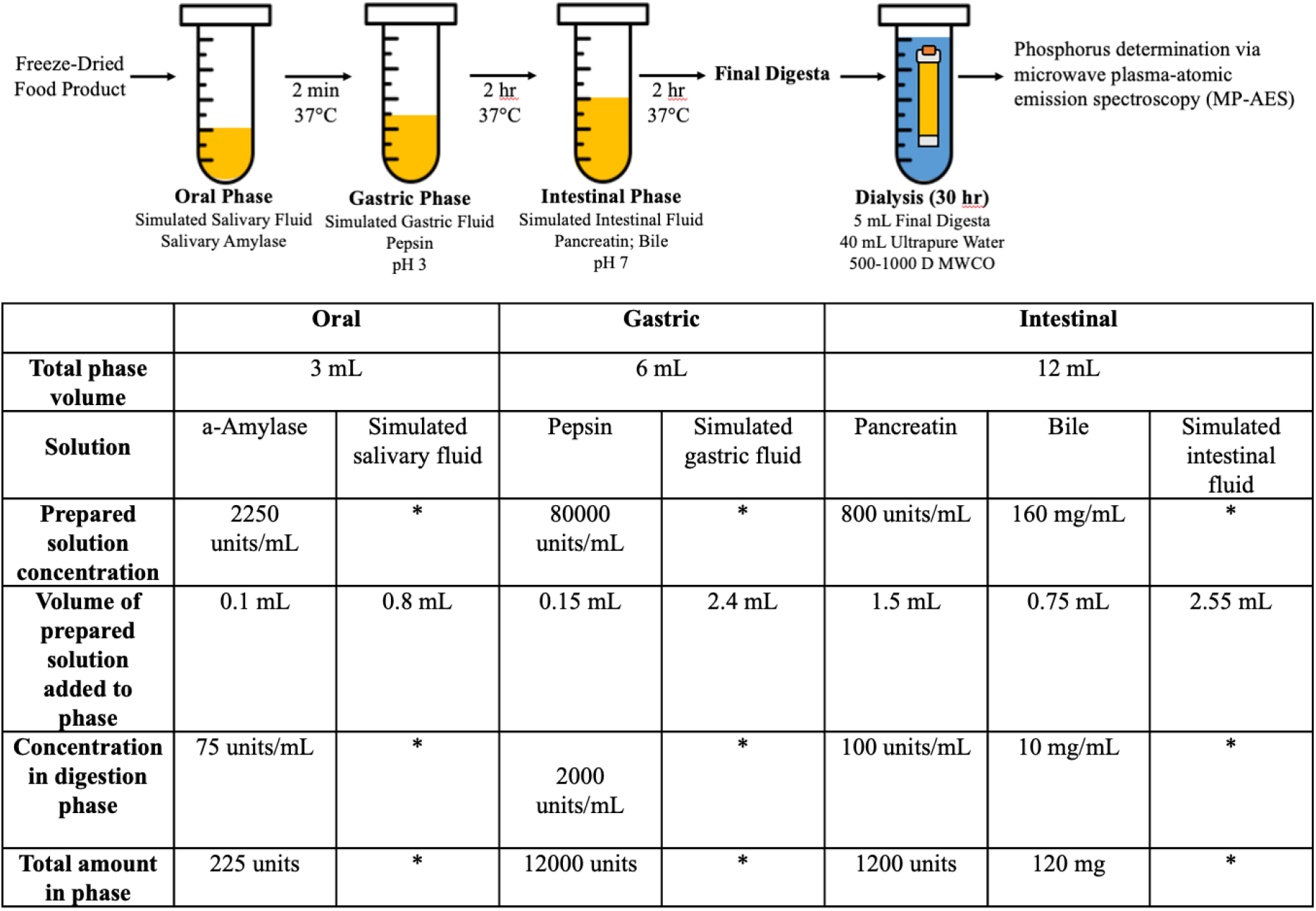
Determination of BioP using an *in vitro* digestion experiment and subsequent dialysis. Freeze-dried food samples were put through an *in vitro* digestion experiment consisting of an oral, gastric, and intestinal phase to simulate human gastrointestinal digestion. Final digesta were subsequently dialyzed against ultrapure water for 30-hours to achieve equilibrium. Final digesta and dialysate samples were analyzed for phosphorus content via microwave plasma atomic emission spectroscopy (MP-AES).

Protein content of each product was determined in duplicate by the Dumas method (AOAC 990.03) using a LECO® FP828 nitrogen analyzer (LECO, St. Joseph, MI, USA). A conversion factor of 6.25 was used for all food products evaluated.

Descriptive statistics were used to characterize data among protein sources (soy, pulse, and animal) and within each food category (beef, dairy, sausage/bacon, chicken/turkey, natural and traditional forms) using Microsoft® Excel® (Version 2303). All other statistical analyses were performed using Statistical Analysis Software (SAS) (Version 9.4 (SAS Institute, Cary NC, USA). Due to the censored data for %PB, a tobit regression analysis was performed to determine the association between %PB and BioP. The strength of the relationship was determined by Pearson correlation. A p-value of <0.05 was considered statistically significant. Mean ± standard deviation of biological replicates are shown.

## Results

### BioP and BioP-to-protein ratio

Average %PB was lowest in pulse protein beef alternatives (32 ± 8%, n=5), followed by soy protein beef alternatives (52 ± 11%, n=4) and was highest in animal protein beef products (89 ± 15%, n=2) (**Supplementary Table 3**). However, the average BioP was similar between pulse and soy protein beef alternatives (83 ± 19 mg/100g, n=5 and 90 ± 13 mg/100g, n=4) (**Figure 2A, Supplementary Table 3**). Both of which were lower than the average BioP from animal protein beef (147 ± 29 mg/100g, n=2). Average BioP-to-protein ratio was similar for soy- and pulse protein beef alternatives (4 ± 1 mg/g for both, n=5, n=4) and lower than animal protein beef (7 ± 0.3 mg/g, n=2) (**Figure 2B, Supplementary Table 1**).

**Figure 2:**
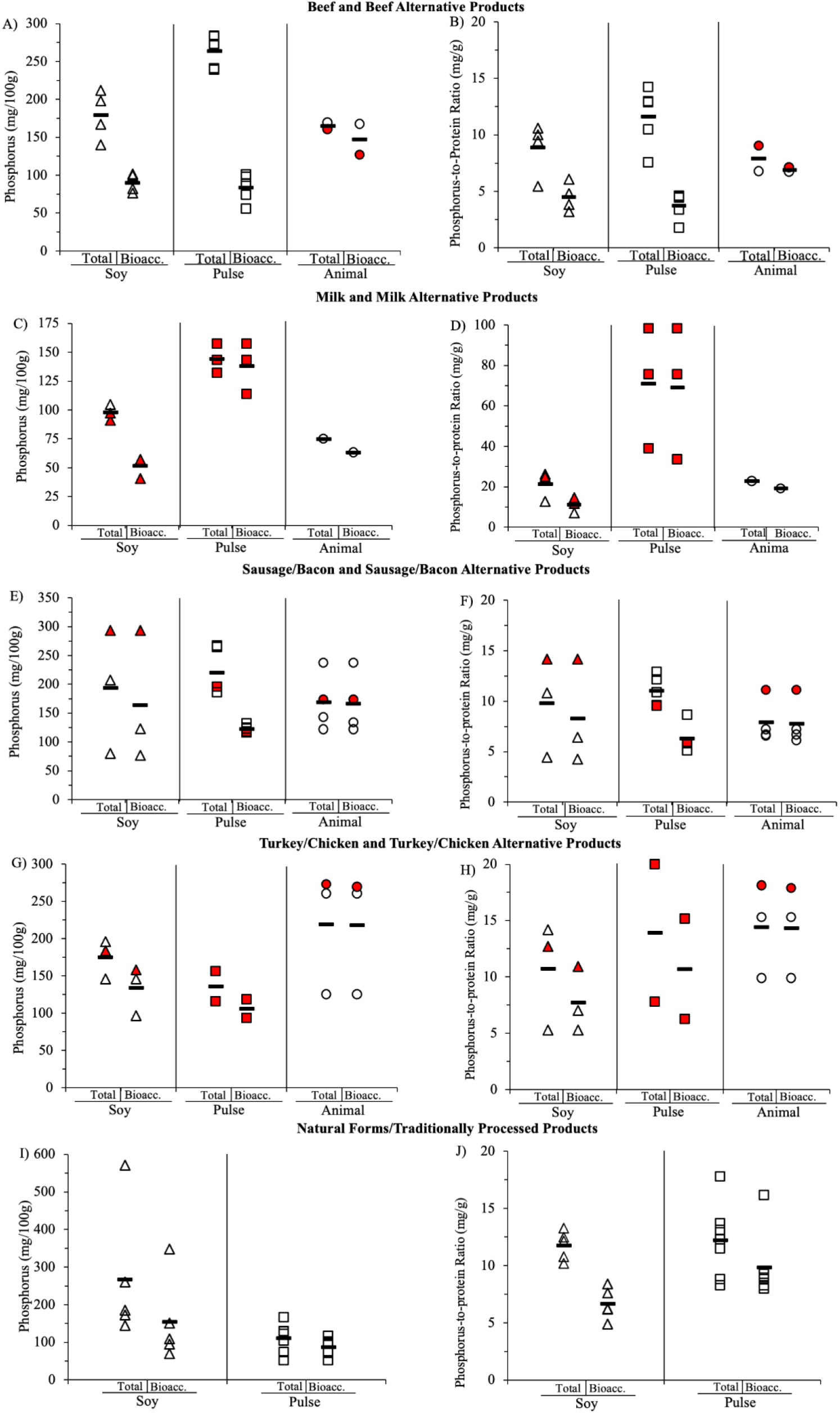
BioP and BioP-to-protein ratio. **A)** Average BioP was similar between soy and pulse-protein beef alternative products. **B)** Average BioP-to-protein ratio was similar for soy and pulse protein beef alternative products, which was lower than the average BioP-to-protein ratio for animal protein beef. **C)** Soy protein milk alternative products and cow’s milk had comparable average BioP. **D)** Average BioP-to-protein ratio was highest in pulse protein and lowest in soy protein milk alternative products. **E)** Average BioP was lower in pulse protein sausage/bacon alternative products compared with soy protein and animal products. **F)** Average BioP-to-protein ratio was lowest in pulse protein sausage/bacon alternative products. **G)** Average BioP was lowest in pulse protein followed by soy protein and highest in animal protein chicken/turkey alternatives and chicken/turkey products. **H)** Average BioP-to-protein ratio was lowest in soy protein followed by pulse protein and highest in animal protein chicken/turkey alternatives and chicken/turkey products. **I)** Average BioP was lower in natural forms/traditionally processed pulse products than soy. **J)** Average BioP-to-protein ratio was lower in natural forms/traditionally processed soy products but was comparable to pulse products. Mean values are shown with a horizontal black line. Triangles represent soy protein products, squares represent pulse protein products, and circles represent animal protein products. Symbols colored red indicate that the product contains at least one inorganic P additive listed on the ingredient list. Bioacc. = bioaccessible. Beef and beef alternatives: Soy n=4, pulse n=5, animal n=2; Milk and milk alternatives: Soy n=3, pulse n=3, animal n=1; Sausage/bacon and sausage/bacon alternatives: Soy n=3, pulse n=5, animal n=4; Chicken/turkey and chicken/turkey alternatives: Soy n=3, pulse n=2, animal n=3; Natural forms/traditionally processed products: Soy n=5, pulse n=7.

Average %PB was higher in pulse protein milk alternatives (95 ± 8%, n=3) than animal (cow’s) milk (85%, n=1) and soy protein milk alternatives (53 ± 7%, n=3) (**Supplementary Table 3**). Average BioP in soy protein milk alternatives (52 ± 9 mg/100g, n=3) was lower but comparable to cow’s milk (63 mg/100g, n=1) (**Figure 2C, Supplementary Table 3**). Notably, average BioP of pulse protein milk alternatives (138 ± 22 mg/100g, n=3) was ∼2x higher than soy protein milk alternatives and cow’s milk (**Figure 2C Supplementary Table 3**). Average BioP-to-protein ratio was lowest in soy protein milk alternatives (11 ± 4 mg/g, n=3), followed by cow’s milk (19 mg/g, n=1) and highest in pulse protein milk alternatives (69 ± 33 mg/g, n=3) (**Figure 2D, Supplementary Table 1**).

Average %PB was lowest in pulse protein sausage alternatives (57 ± 11%, n=5), followed by soy protein sausage/bacon alternatives (85 ± 22%, n=3) and was highest in animal protein sausage/bacon products (98 ± 3%, n=4) (**Supplementary Table 3**). Pulse protein sausage alternatives had the lowest average BioP (123 ± 6 mg/100g, n=5), whereas BioP for soy protein sausage/bacon alternatives and animal protein sausage/bacon products were similar (164 ± 114 mg/100g, n=3 and 166 ± 52 mg/100g, n=4 respectively) (**Figure 2E, Supplementary Table 3**). Average BioP-to-protein ratio was lowest in pulse protein sausage alternatives (6 ± 1 mg/g, n=5) and was comparable between soy protein sausage/bacon alternatives and animal protein sausage/bacon products (8 ± 5 mg/g, n=3 and 8 ± 2 mg/g, n=4) (**Figure 2F, Supplementary Table 3**).

Average %PB was comparable for pulse protein and soy protein chicken/turkey alternatives (78 ± 3%, n=2 and 78 ± 26%, n=3) while animal protein chicken/turkey had the highest average %PB (100 ± 1%, n=3) (**Supplementary Table 3**). Average BioP was 106 ± 18 mg/100g (n=2) for pulse protein chicken/turkey alternatives, ∼1.3x and 2x lower than soy protein chicken/turkey alternatives and animal protein chicken/turkey products (134 ± 33 mg/100g, n=3 and 218 ± 81 mg/100g, n=3) (**Figure 2G, Supplementary Table 3)**. Average BioP-to-protein ratio was lowest in soy protein chicken/turkey alternatives (8 ± 3 mg/g, n=3) followed by pulse protein chicken/turkey alternatives (11 ± 6 mg/g, n=2) and was highest in animal protein chicken/turkey products (14 ± 4 mg/g, n=3) (**Figure 2H, Supplementary Table 3**).

Average %PB was lower in natural forms/traditionally processed soy products (i.e., tofu, tempeh, soybeans etc.) compared with natural forms/traditionally processed pulse products (i.e., chickpeas, green lentils, fava beans etc.) (56 ± 7%, n=5 and 82 ± 15%, n=7, respectively) (**Supplementary Table 3**). Interestingly, average BioP was lower in natural forms/traditionally processed pulse products compared with natural forms/traditionally processed soy products (87 ± 22 mg/100g, n=7 and 154 ± 112 mg/100g, n=5) (**Figure 2I**). Canned pulse products had considerably higher average %PB compared to the same prepared dry pulse products (100 ± 0%, n=2 and 64 ± 5%, n=2) but average BioP was lower in the canned products (63 ± 15 mg/100g, n=2 and 86 ± 22 mg/100g, n=2). Natural forms/traditionally processed soy products had a lower but comparable average BioP-to-protein ratio to natural forms/traditionally processed pulse products (7 ± 1 mg/g, n=5 and 10 ± 3 mg/g, n=7) (**Figure 2J, Supplementary Table 3**).

### Relationship between %PB and BioP and the presence of inorganic P additives

%PB and BioP were significantly associated (Wald X^2^ = 9.34, p=0.002) with a moderate strength of relationship (r=0.46, p=0.0002) (**Figure 3)**. Of all 63 products evaluated, 15 listed at least one inorganic P additive on the ingredient list. Within each food category, the presence of ≥1 inorganic P additive on the ingredient list resulted in an average %PB of 78-87% which overlaps with the %PB of products without an inorganic P additive on the ingredient list (average range: 46-87%). Notably, the average %PB of soy protein milk alternative products with and without additives were similar (52 ± 10%, n=2 and 54%, n=1, respectively) (**Supplementary Table 3**).

**Figure 3:**
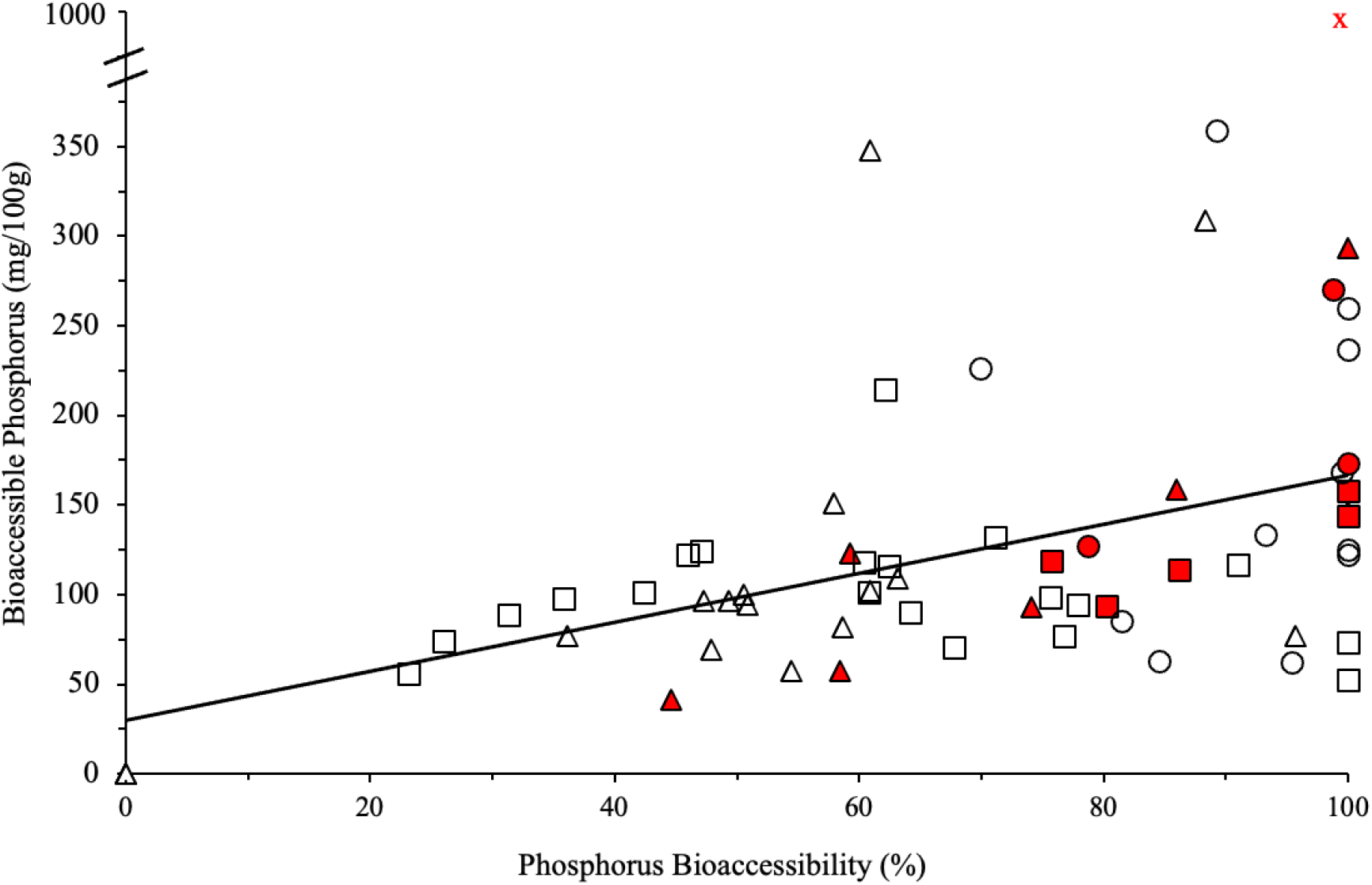
Relationship between %PB and BioP. A moderate strength relationship was observed for %PB and BioP. Triangles represent soy protein products, squares represent pulse protein products, and circles represent animal protein products. Symbols colored red indicate that the product contains at least one inorganic P additive listed on the ingredient list. Outliers are shown with a red “x”, n=62 and n=1 outlier (total n=63).

## Discussion

The bioavailability approach to dietary phosphorus management in CKD has gained much interest and support over recent years, where plant sources are generally emphasized over animal sources, and inorganic P additives are to be avoided.^1,2,12,25^ However, there is uncertainty surrounding advice on processed plant protein products, due to possible effects of various processing methods on phosphorus bioaccessibility as well as the possible inclusion of inorganic P additives in these products.^3,4,22,26,27^ Overall, we found that most of the processed plant protein products evaluated in this pilot study had *lower or similar* phosphorus bioaccessibility compared with animal protein counterparts, which aligns with previous findings that plant sources generally have lower phosphorus bioaccessibility than animal sources.^3,6,15^ However, we also found wide ranges of %PB and BioP, and some of the processed plant protein products had substantially more BioP than animal protein counterparts (e.g., pulse protein milk alternatives vs. cow’s milk or soy protein milk alternative products).

It is critical for clinicians making dietary recommendations for people with CKD to understand the relationship between total phosphorus content, % phosphorus bioaccessibility and bioavailability, and amount of bioaccessible or bioavailable phosphorus in foods. We propose using consistent and specific terminology for “bioaccessibility” vs. “bioavailability” (as shown in **Box 1**) to support clear communication in discussions of these concepts within the field. This is important because conflation of various terms when referring broadly to phosphorus “bioavailability” has contributed to some misinterpretations of practical significance. The first example of this is that discussions on this topic usually refer to (estimated) differences in % phosphorus bioavailability by source but based primarily on data describing %PB (not bioavailability), which has led to these estimates being misinterpreted as % phosphorus absorption. For example, it is commonly misstated that ∼100% of phosphorus from inorganic P additives *is absorbed* (rather than just in accessible form). But data suggest that only ∼73% of phosphorus from inorganic phosphate is absorbed in healthy adults.^28^ The second misinterpretation occurs when a focus on %PB disregards total phosphorus content and thus the actual amount of BioP. For example, in this study, canned chickpeas had 100%PB, but because total phosphorus was low, BioP load was low. Ideally, %PB and the total phosphorus content should always be considered together to determine BioP load (mg) as the measure with the most practical importance. This approach would apply a %PB factor to total phosphorus content to estimate BioP load for foods/beverages (e.g., mg per serving, per 100 g, or per 100 kcal).^12^ Development of a valid and reliable predictive equation that estimates BioP for a variety of foods/beverages is needed, but the scarcity of data currently precludes this.

Processed foods are generally discouraged based on the possible effects of processing techniques, and the widespread use of inorganic P additives in many processed foods. This rationale has been applied to processed plant protein products but based on limited evidence. We observed a wide range in %PB in processed plant protein products (∼32-100%) which overlaps, but extends higher than, %PB ranges previously reported for plant sources (∼10-50%).^3,6,15^ Phytic acid content, methods of processing and formulation, and food matrices may affect %PB and contribute to the wide range observed.^29–35^ For example, phytic acid content can be reduced by dehulling, germination, extrusion, soaking, boiling, and microwaving.^36–40^ Indeed, average %PB of cooked, natural or traditionally-processed (e.g., tofu) forms of legumes in this pilot study were higher than previously reported for similar uncooked/raw legumes.^15^

Processed foods may contain inorganic P additives,^41–43^ which can have up to 100% phosphorus bioaccessiblity.^44,45^ Evidence suggests that widespread use of inorganic P additives contributes substantially to average dietary intake,^42,46^ and reducing intake of these additives has been shown effective for reducing serum phosphorus in adults receiving hemodialysis without adversely affecting overall nutritional status.^47,48^ However, for individual food products, the amount of added phosphorus from inorganic P additives are unknown and could have negligible to substantial effects on the BioP load. Our pilot data indicate that the presence of inorganic P additives on the ingredient list is not associated with either high %PB nor BioP load of the product. Notably, the soy protein milk alternatives had similar BioP compared with the cow’s milk comparator, but the pulse protein milk alternatives had ∼2x higher BioP than the soy protein and cow’s milk products. This was despite inorganic P additives being listed for all the pulse protein milk alternatives and all but one of the soy protein milk alternatives. When protein content was considered, the soy protein milk alternatives had ∼1.7x *lower* BioP-to-protein ratio compared with the cow’s milk. These data suggest that soy protein milk alternatives could be a favorable choice for people with CKD, particularly those receiving dialysis who are advised to consume higher protein while limiting dietary phosphorus.^49,50^

A strength of the current study was that the *in vitro* digestion experiments were based on the standardized INFOGEST protocol, internationally developed to simulate adult human digestion in a static *in vitro* system.^24^ But, *in vitro* systems may not reflect *in vivo* bioaccessibility nor bioavailability. However, these data provide valuable information under standardized conditions that can generate hypotheses for future research. The wide variety of foods evaluated in this pilot study was a strength, but a corresponding limitation was the small number of products evaluated per category (e.g., milk, beef) and one sample per product. Broader representation of foods within categories would be needed to reasonably perform inferential statistical analysis for differences among categories.

This pilot study demonstrates that some processed plant protein products may offer similar or lower BioP per serving and per gram of protein compared with some animal protein comparator products, including some products that list inorganic P additive ingredients. Despite these preliminary data, the current broad guidance to avoid more highly processed foods and foods that list inorganic P additive ingredients remains a responsible approach until there is more information on phosphorus bioaccessibility on a wider range of products, as well as mandatory labeling of total and added inorganic P content on the Nutrition Facts label.^51^ However, our data should provide incentive for manufacturers to consider voluntarily reporting total and phosphorus content for plant protein products with favorable profiles. Overall, this pilot study suggests that the favorable attributes of plant protein sources for lower dietary P burden are not solely limited to “natural”, unprocessed sources.

## Acknowledgements

The authors would like to thank the undergraduate student workers in the Hill Gallant Laboratory, Heily Yin and Lauren Palaia, for their assistance with performing the experiments. Additionally, the authors thank Dr. Pam Ismail and Nigel Kang for their assistance in determining the protein content of the food products evaluated. Portions of this work have been presented at the National Kidney Foundation Spring Clinical Meeting 2022 (P378, P381) and 2023 (P390), American Society of Nephrology Kidney Week 2022 (TH-PO801) and in the doctoral dissertation of KMB: Burstad, Kendal. (2023) Dietary Phosphorus in Chronic Kidney Disease: Effects of Amount, Source and Bioaccessibility on Intestinal Absorption and Health Outcomes. Retrieved from the University of Minnesota Digital Conservancy, https://hdl.handle.net/11299/257023.

## Box 1.

**Terminology for Phosphorus Bioaccessibility and Bioavailability**

### Bioaccessibility Terms

***Percent Phosphorus (% P) bioaccessibility***: the *percent of total* phosphorus that *is accessible* for absorption.

***Bioaccessible phosphorus (P):*** the *absolute amount (mg)* of phosphorus that *is accessible* for absorption.

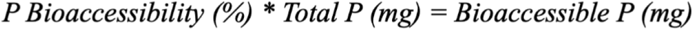

### Bioavailability Terms

***Percent Phosphorus (%P) bioavailability***: the *percent of total* phosphorus that *is absorbed* and available for utilization by the body.

***Bioavailable phosphorus (P):*** the *absolute amount (mg)* of phosphorus that *is absorbed* and available for utilization by the body.

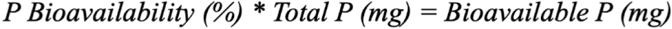

