## Supplemental File for "Phosphorus Bioaccessibility of Processed Soy and Pulse Protein Products Measured by *In Vitro* Simulation of Human Digestion"

**Supplementary Methods:**

Product Procurement and Preparation

Pulse protein, soy protein, and animal protein counterpart products were selected for this study based on popular demand in the United States using retail sales data from the Good Foods Institute and SPINS data[1] and by sorting products online using "best seller" filters. All products were purchased from nationwide retailers located in the Minneapolis-St. Paul, Minnesota metro area in the United States between August 2021-January 2022. Natural forms/traditionally processed plant protein products (e.g., tofu, peanut butter) (n=17) were also evaluated for comparison. Information on inorganic P-containing food additives was obtained from the ingredient list on each product package **(Supplementary Table 1)**.

Products were prepared according to package directions which included methods such as cooking in a microwave, oven, or on a skillet. Dried legume products were prepared by soaking then subsequent boiling (**Supplementary Table 2**). Tap water was used in the cooking process when necessary. No additional oils, spices, or ingredients were used. Prepared products were frozen at -20°C for at least 24 hours then freeze dried (FTS systems, Dura-Dry Condense Module Bulk Tray Dryer) for 4-7 days. Following freeze drying, products were pulverized into small particles (~2.2mm or less) using a mallet and mortar and pestle. All freeze-dried samples were kept sealed at room temperature until use.

Measurement of Total Phosphorus

Five replicates of each freeze-dried food sample were dry-ashed in a muffle furnace (Thermo Scientific, Thermolyne Furnace, Atmosphere Controlled Ashing, Model F30400). Briefly, 2-3g of freeze-dried food sample were weighed into porcelain crucibles and placed in a muffle furnace. Samples were heated to 300°C at a rate of 3°C per minute and held at 300°C for 16 hours, then ramped to 600°C at a rate of 3°C per minute and held at 600°C for 72 hours. Ashed sample was dissolved with 1 mL of concentrated (70%) nitric acid and diluted with ultrapure water to 25 mL. Samples were filtered using a 0.45µm filter to remove particulates and then analyzed by microwave plasma atomic emission spectroscopy (MP-AES) (Agilent Technologies, model 4210) using 213.618 nm for emission detection of phosphorus. Total calcium, magnesium, potassium and sodium were also determined for the freeze-dried food samples (pre-digestion) as described above for phosphorus and are provided in **Supplementary Table 4**.

In Vitro Digestion

Simulated salivary, gastric and intestinal fluids and enzyme solutions were prepared to closely mimic the composition of human digestive fluids according to the INFOGEST protocol recommendations.[2] Simulated salivary, gastric and intestinal fluids consisted of the following electrolyte solutions: 0.5M potassium chloride, 0.5M monopotassium phosphate, 1M sodium bicarbonate, 2M sodium chloride, 0.15M magnesium chloride hexahydrate, 0.5M ammonium carbonate, and 6M hydrochloric acid with the exception that the simulated salivary fluid did not include sodium chloride and the simulated intestinal fluid did not include ammonium carbonate. Stock solutions of each simulated fluid were prepared at a 1.25x concentration and were kept at -20°C until use. Prior to use in the *in vitro* digestions, simulated fluids were prewarmed to 37°C in a shaking water bath.

All enzyme solutions were prepared the day of the *in vitro* digestions. α-amylase (from *bacillus sp.,* Sigma Aldrich A6380) and pepsin (from porcine gastric mucosa, Sigma Aldrich P7012) solutions were prepared with ultrapure water, whereas pancreatin (from porcine pancreas, Sigma Aldrich P7545) and bile (porcine, Sigma Aldrich B8631) solutions were prepared with simulated intestinal fluid. The concentration of α-amylase solution in the oral phase was 75 units/mL and pepsin solution in the gastric phase was 2000 units/mL, determined according to the manufacturer certificate of analysis. The concentration of bile in the intestinal phase was 10 mg/mL and the concentration of pancreatin solution in the intestinal phase was normalized to the amount of trypsin activity, 100 units/mL (**Figure 1**). The trypsin activity of pancreatin was determined based on the method described in the INFOGEST protocol.[2] Briefly, 2.6 mL of a 46 mM Tris/HCl buffer and 0.3 mL of 10 mM na-tosyl-arginine-methyl-ester were pipetted into a cuvette and warmed to 25°C. Pancreatin was added to the cuvette in varying concentrations 0.1, 0.5, 0.75, 1 and 1.5 mg/mL, and the absorbance increase at 247 nm was measured by UV-vis spectrometry and recorded in continuum over 10 minutes, until levelling off.

Each freeze-dried sample was digested in triplicate. In addition, a 'process blank' sample, using ultrapure water, was prepared to determine background phosphorus. Each food product was pre-tested to determine adjustments needed by HCl and NaOH to achieve target pH levels of the gastric (pH= 3, range 2.8-3.2) and intestinal (pH= 7, range 6.8-7.2) simulated digestion phases. For the oral phase, one gram of freeze-dried food sample or ultrapure water (blank) was weighed into a 50 mL centrifuge tube. Then 0.8 mL simulated salivary fluid, 0.1 mL α-amylase, 0.005 mL 0.3 M calcium chloride ($CaCl_2$), and 1.095 mL ultrapure water were added to each tube. The final volume of the oral phase was 3 mL. Then, the tubes were placed in a 37°C shaking water bath and oscillated at 110-115 oscillations per minute (OPM) for 2 minutes. For the gastric phase, 2.4 mL simulated gastric fluid, 0.15 mL pepsin, and 0.00015 mL 0.3 M $CaCl_2$ were added to each tube, and pH adjusted to 3. Ultrapure water was added to achieve a final gastric phase volume of 6 mL. Tubes were then returned to the shaking water bath (37°C and 110-115 OPM) for 2 hours. After 2 hours, 2.55 mL simulated intestinal fluid, 1.5 mL pancreatin, 0.75 mL bile, and 0.012 mL 0.3 M $CaCl_2$ were added to start the intestinal phase and the pH was adjusted to 7. Ultrapure water was added to reach a final intestinal phase volume of 12 mL. The tubes were again incubated in the shaking water bath (37°C and 110-115 OPM) for 2 hours. Directly following the intestinal phase, the final digesta was diluted with ultrapure water to 25 mL.

<u>pH Testing</u>

In preparation for the *in vitro* digestion experiments, all products underwent pH testing in which the amount of 5M HCl and 5M NaOH required to achieve target pH levels of the gastric (pH=3, range 2.8-3.2) and intestinal (pH=7, range 6.8-7.2) simulated digestion phases were determined. Briefly, products were put through a test *in vitro* digestion using the protocol described above, except digestive enzymes were not utilized. After adding the required reagents to the gastric phase, the pH was tested and adjusted with known volumes of 5 M HCl or 5 M NaOH until reaching a pH of 3 (range 2.8-3.2). This was repeated for the intestinal phase until a pH of 7 (range 6.8-7.2) was reached. The amount of volume required to achieve the desired pH at each phase was recorded and utilized when performing the *in vitro* digestions.

Bioaccessible Phosphorus Determination

Following dialysis, dialysate and diluted digesta were analyzed for phosphorus content using MP-AES. Diluted digesta of each sample were dry-ashed to determine total phosphorus content after the *in vitro* digestion process. Briefly, 2-3g of diluted digesta were pipetted into porcelain crucibles and placed in a muffle furnace. Samples were heated to 60°C at a rate of 3°C per minute and held at 60°C for 24 hours, then ramped to 600°C at a rate of 3°C per minute and held at 600°C for 24 hours. Dry-ashed samples were dissolved in 1 mL of concentrated (70%) nitric acid and diluted to 25 mL with ultrapure water (final nitric acid concentration 2.8%).

Bioaccessible Phosphorus Calculation

Ashed diluted digesta and dialysate of each sample were run on the MP-AES for determination of phosphorus on the same days with the same calibration curve and analytical blanks. Phosphorus of analytical blanks was averaged and subtracted from the experimental blank to accurately determine the contribution of background phosphorus from the *in vitro* digestion experiments. The contribution of background phosphorus was then subtracted from the phosphorus of the ashed diluted digesta samples to determine total phosphorus post-digestion. Contribution of background phosphorus was found to be negligible in the dialysate. The phosphorus values from the post-digestion and dialysate were used to calculate percent bioaccessible phosphorus. The percent bioaccessible phosphorus was then applied back to the total phosphorus of the whole food product to calculate the bioaccesible phosphorus.

[1] Retail sales data: Plant-based meat, eggs, dairy. Good Foods Institute. Accessed May 8, 2021. https://gfi.org/resource/marketresearch/

[2] Brodkorb A, Egger L, Alminger M, et al. INFOGEST static in vitro simulation of gastrointestinal food digestion. *Nat Protoc*. 2019;14:24. doi:10.1038/s41596-018-0119-1

**Supplementary Table 1. Product details**

| Food Category | Protein Category | Product | Listed Inorganic Phosphate Additives |
|---|---|---|---|
| Beef | Soy | Impossible - Plant-Based Burger Ground | N/A |
| | | Incogmeato MorningStar Farms - Burger Patties | N/A |
| | | Gardein - Plant-Based Ground Be'f | N/A |
| | | MorningStar Farms - Vegan Meat Lovers (Patties) | N/A |
| | Pulse | Beyond Meat - Beyond Beef Beefy Crumbles | N/A |
| | | Sweet Earth - Awesome Grounds | N/A |
| | | Beyond Meat - Beyond Beef Plant Based Patties | N/A |
| | | Gardein - Ultimate Plant-Based Burger | N/A |
| | | Dr. Praeger's - All American Plant-Based Burgers | N/A |
| | Animal | Bubba Burger - Original Burger | N/A |
| | | BallPark - Beef Patty | sodium phosphate |
| Milk | Soy | Silk - Original Soymilk | tricalcium phosphate |
| | | Pacific Foods - Ultra Soy Plant-Based Beverage Original | tricalcium phosphate |
| | | Silk - Ultra Original Soy Protein Beverage | N/A |
| | Pulse | Ripple - Unsweetened Plant-Based Milk Original | tricalcium phosphate; dipotassium phosphate |
| | | Sproud - Plant-Based Milk Original | dipotassium phosphate |
| | | Not Milk - Plant-Based Milk Alternative 2% Reduced Fat | dipotassium phosphate; monocalcium phosphate |
| | Animal | Kemps - 2% Reduced Fat Milk | N/A |
| Yogurt & Cheese | Soy | Silk - Dairy-Free Yogurt Vanilla | tricalcium phosphate; dipotassium phosphate |
| | | Field Roast - Vegan Chao Shreds Creamy Original | N/A |
| | | Field Roast - Vegan Chao Slices Creamy Original | N/A |
| | Pulse | Miyoko's - Cultured Vegan Cheese Farmhouse Cheddar | N/A |
| | | Daiya - Cheddar Style Slices | N/A |

| | | | |
|---|---|---|---|
| | Animal | Yoplait - Original French Vanilla | N/A |
| | | Kraft - Sharp Cheddar Shredded Cheese | N/A |
| | | Kraft Singles - American | calcium phosphate; sodium phosphate |
| Sausage & Bacon | Soy | Gardein - Plant-Based Sliced Italian Saus'ge | N/A |
| | | MorningStar Farms - Veggie Sausage Links | N/A |
| | | MorningStar Farms - Veggie Bacon Strips | sodium phosphate; monocalcium phosphate; sodium tripolyphosphate |
| | Pulse | Beyond Meat - Beyond Sausage Plant-Based Links Brat Original | N/A |
| | | Raised & Rooted - Bratwurst Plant Based Sausage | tetrasodium pyrophosphate |
| | | Beyond Meat - Beyond Sausage Plant-Based Links Hot Italian | N/A |
| | | Beyond Meat - Beyond Breakfast Sausage Plant-Based Links Classic | N/A |
| | | Lightlife - Plant-Based Breakfast Patties | N/A |
| | Animal | Johnsonville - Original Bratwurst | N/A |
| | | Johnsonville - Italian Sausage Mild | N/A |
| | | Jimmy Dean - Original Pork Sausage Patties | sodium phosphate |
| | | Hormel - Black Label Original Bacon | N/A |
| Chicken & Turkey | Soy | Wholesome Provisions - Just Like Chicken Textured Vegetable Protein | N/A |
| | | MorningStar Farms - Original Chik Patties | sodium acid pyrophosphate |
| | | Tofurky - Plant Based Deli Slices Oven Roasted | N/A |
| | Pulse | Raised & Rooted - Plant Based Nuggets | sodium acid pyrophosphate |
| | | Beyond Meat - Beyond Chicken Plant Based Breaded Tenders | sodium acid pyrophosphate; monocalcium phosphate; sodium phosphates |
| | Animal | Tyson - Boneless Skinless Chicken Breasts | N/A |

| | | | |
|---|---|---|---|
| | | Tyson - Chicken Patties | N/A |
| | | Hillshire Farm - Oven Roasted Turkey Breast | sodium phosphate |
| Natural Forms & Traditional Processed Plant-Based Protein Products | Soy | Tofu | N/A |
| | | LightLife Tempeh | N/A |
| | | Bird's Eye Edamame | N/A |
| | | Seapoint Farms Edamame Dry Roasted | N/A |
| | | Soymerica Soybeans | N/A |
| | Pulses | Chickpeas (canned) | N/A |
| | | Chickpeas (Dry) | N/A |
| | | Green Lentils (Dry) | N/A |
| | | Yellow Split Peas (Dry) | N/A |
| | | Fava Bean (Canned) | N/A |
| | | Fava Bean (Dry) | N/A |
| | | Mung Bean (Dry) | N/A |
| | Peanut | Jif Peanut Butter (creamy) | N/A |
| | Soy | WowButter Soy Spread (creamy) | N/A |
| | Wheat | Barilla Pasta | N/A |
| | Soy | Simply Nature Soybean Spaghetti | N/A |
| | Pulse | Banza Spaghetti | N/A |

**Supplementary Table 1. Product details.** Product details including food category, protein category, specific food product evaluated, and the presence of inorganic phosphate-containing foods additives found on the ingredient list for each product.

**Supplementary Table 2. Product cooking methods**

| Food Category | Protein Category | Product | Cooking Mode | Cooking Details |
|---|---|---|---|---|
| Beef | Soy | Impossible - Plant-Based Burger Ground | Skillet | Med-high heat; 8 min; from thawed |
| | | Incogmeato MorningStar Farms - Burger Patties | Skillet | Med-high heat; 4 min per side; from frozen |
| | | Gardein - Plant-Based Ground Be'f | Skillet | Med-high heat; 15 min; from frozen |
| | | MorningStar Farms - Vegan Meat Lovers (Patties) | Skillet | Med-high heat; 18 min; from frozen |
| | Pulse | Beyond Meat - Beyond Beef Beefy Crumbles | Skillet | Med heat; 6 min; from frozen |
| | | Sweet Earth - Awesome Grounds | Skillet | Med-high heat; 13 min; from frozen |
| | | Beyond Meat - Beyond Beef Plant Based Patties | Skillet | Med-high heat; 10 min; from thawed |
| | | Gardein - Ultimate Plant-Based Burger | Skillet | Med heat; 6-7 min per side; from frozen |
| | | Dr. Praeger's - All American Plant-Based Burgers | Skillet | Med heat; 10 min per side; from frozen |
| | Animal | Bubba Burger - Original Burger | Skillet | Med heat; 7-10 min per side; from frozen |
| | | BallPark - Beef Patty | Skillet | Med heat; 8-10 min; from frozen |
| Milk | Soy | Silk - Original Soymilk | N/A | |
| | | Pacific Foods - Ultra Soy Plant-Based Beverage Original | N/A | |
| | | Silk - Ultra Original Soy Protein Beverage | N/A | |
| | Pulse | Ripple - Unsweetened Plant-Based Milk Original | N/A | |
| | | Sproud - Plant-Based Milk Original | N/A | |
| | | Not Milk - Plant-Based Milk Alternative 2% Reduced Fat | N/A | |
| | Animal | Kemps - 2% Reduced Fat Milk | N/A | |
| | Soy | Silk - Dairy-Free Yogurt Vanilla | N/A | |

| | | | | |
|---|---|---|---|---|
| Yogurt & Cheese | | Field Roast - Vegan Chao Shreds Creamy Original | N/A | |
| | | Field Roast - Vegan Chao Slices Creamy Original | N/A | |
| | Pulse | Miyoko's - Cultured Vegan Cheese Farmhouse Cheddar | N/A | |
| | | Daiya - Cheddar Style Slices | N/A | |
| | Animal | Yoplait - Original French Vanilla | N/A | |
| | | Kraft - Sharp Cheddar Shredded Cheese | N/A | |
| | | Kraft Singles - American | N/A | |
| Sausage & Bacon | Soy | Gardein - Plant-Based Sliced Italian Saus'ge | Skillet | Med-high heat; 8 min |
| | | MorningStar Farms - Veggie Sausage Links | Skillet | Med heat; 10 min |
| | | MorningStar Farms - Veggie Bacon Strips | Skillet | Med heat; 10 min |
| | Pulse | Beyond Meat - Beyond Sausage Plant-Based Links Brat Original | Skillet | Med-high heat; 12 min |
| | | Raised & Rooted - Bratwurst Plant Based Sausage | Skillet | Med heat; 10 min |
| | | Beyond Meat - Beyond Sausage Plant-Based Links Hot Italian | Skillet | Med heat; 9 min |
| | | Beyond Meat - Beyond Breakfast Sausage Plant-Based Links Classic | Skillet | Med heat; 10 min |
| | | Lightlife - Plant-Based Breakfast Patties | Skillet | Med heat; 12 min |
| | Animal | Johnsonville - Original Bratwurst | Skillet | Med-high heat for 5 min; then added water and covered for 20 min |
| | | Johnsonville - Italian Sausage Mild | Skillet | Med-high heat; 11 min |
| | | Jimmy Dean - Original Pork Sausage Patties | Skillet | Med heat; 7 min |
| | | Hormel - Black Label Original Bacon | Skillet | Med heat; 18-22 min |
| Chicken & Turkey | Soy | Wholesome Provisions - Just Like Chicken Textured Vegetable Protein | Skillet | Med-high heat; 7 min; 1:1 ratio of (chicken: water) |

| | | | | |
|---|---|---|---|---|
| | | MorningStar Farms - Original Chik Patties | Oven | 375 degrees F; 16-18 min |
| | | Tofurky - Plant Based Deli Slices Oven Roasted | N/A | |
| | Pulse | Raised & Rooted - Plant Based Nuggets | Oven | 400 degrees F; 10 min |
| | | Beyond Meat - Beyond Chicken Plant Based Breaded Tenders | Oven | 425 degrees F; 4 min per side |
| | Animal | Tyson - Boneless Skinless Chicken Breasts | Oven | 350 degrees F; 60 min |
| | | Tyson - Chicken Patties | Oven | 400 degrees F; 17-20 min |
| | | Hillshire Farm - Oven Roasted Turkey Breast | N/A | |
| Natural Forms & Traditional Processed Plant-Based Protein Products | Soy | Tofu | Skillet | Med-high heat; 6 min (cubed) |
| | | LightLife Tempeh | Skillet | Med-high heat; 8 min |
| | | Bird's Eye Edamame | Microwave | Heat 3.5 min; let stand 1 min |
| | | Seapoint Farms Edamame Dry Roasted | N/A | |
| | | Soymerica Soybeans | Boil | Soaked in a 1:3 (bean:water) ratio overnight; Drained; Rinsed; Added to boiling water at same ratio for 3 hours |
| | Pulses | Chickpeas (canned) | Rinse | Rinsed with water |
| | | Chickpeas (Dry) | Boil | Soaked in a 1:3 (pea:water) ratio overnight; Drained; Rinsed; Added to boiling water at same ratio for 1.5 hours |
| | | Green Lentils (Dry) | Boil | Added to boiling water at 1:4 (lentil: water) ratio; simmered 20 min |
| | | Yellow Split Peas (Dry) | Boil | Peas added to water in a 1:2 (pea:water) ratio; brought to boil; simmered 30 min |
| | | Fava Bean (Canned) | Microwave | Drained but not rinsed; heated 3 min |
| | | Fava Bean (Dry) | Boil | Soaked in a 1:10 (bean:water) ratio overnight; drained; peeled; boiled at same ratio for 10 min |
| | | Mung Bean (Dry) | Boil | Soaked in a 1:3 (bean:water) ratio for 15 min; brought to boil in same ratio; simmered 30 min; let stand 10 min |
| | Peanut | Jif Peanut Butter (creamy) | N/A | |

| | | | | |
|---|---|---|---|---|
| | Soy | WowButter Soy Spread (creamy) | N/A | |
| | Wheat | Barilla Pasta | Boil | 10 min |
| | Soy | Simply Nature Soybean Spaghetti | Boil | 4.5 min; 8 cups water used to boil entire package |
| | Pulse | Banza Spaghetti | Boil | 9 min |

**Supplementary Table 2. Product cooking methods.** Cooking mode and details for each evaluated product.

**Supplementary Table 3. Bioaccessible phosphorus and bioaccessible phosphorus-to-protein ratio**

| Food Category | Protein Category | Product | Total P (mg/100g) | Percent Bioaccessible P (%) | Bioaccessible P (mg/100g) | Protein (g/100g) | Serving Size (g) | Total P (mg/serving) | Bioaccessible P (mg/serving) | Total P-to-protein Ratio (mg/g) | Bioaccessible P-to-protein Ratio |
|---|---|---|---|---|---|---|---|---|---|---|---|
| Beef | Soy | Impossible - Plant-Based Burger Ground | 167 | 61 | 102 | 17 | 113 | 189 | 115 | 10 | 6 |
| | | Incogmeato MorningStar Farms - Burger Patties | 212 | 36 | 77 | 20 | 120 | 254 | 92 | 11 | 4 |
| | | Gardein - Plant-Based Ground Be'f | 198 | 51 | 100 | 21 | 87 | 172 | 87 | 9 | 5 |
| | | MorningStar Farms - Vegan Meat Lovers (Patties) | 140 | 59 | 82 | 26 | 113 | 158 | 93 | 5 | 3 |
| | Pulse | Beyond Meat - Beyond Beef Beefy Crumbles | 241 | 23 | 56 | 32 | 55 | 133 | 31 | 8 | 2 |
| | | Sweet Earth - Awesome Grounds | 284 | 26 | 74 | 22 | 85 | 241 | 63 | 13 | 3 |
| | | Beyond Meat - Beyond Beef Plant Based Patties | 273 | 36 | 98 | 21 | 113 | 308 | 110 | 13 | 5 |
| | | Gardein - Ultimate | 239 | 42 | 101 | 23 | 113 | 270 | 114 | 10 | 4 |

| | | | | | | | | | | | |
|---|---|---|---|---|---|---|---|---|---|---|---|
| | | Plant-Based Burger | | | | | | | | | |
| | | Dr. Praeger's - All American Plant-Based Burgers | 283 | 31 | 89 | 20 | 113 | 320 | 100 | 14 | 4 |
| | Animal | Bubba Burger - Original Burger | 169 | 99 | 168 | 25 | 151 | 255 | 254 | 7 | 7 |
| | | [a]BallPark - Beef Patty | 161 | 79 | 127 | 18 | 77 | 124 | 98 | 9 | 7 |
| Milk | Soy | [a]Silk - Original Soymilk | 91 | 45 | 41 | 4 | 240 | 219 | 98 | 26 | 12 |
| | | [a]Pacific Foods - Ultra Soy Plant-Based Beverage Original | 97 | 58 | 57 | 4 | 240 | 233 | 136 | 25 | 15 |
| | | Silk - Ultra Original Soy Protein Beverage | 105 | 54 | 57 | 8 | 240 | 251 | 137 | 13 | 7 |
| | Pulse | [a]Ripple - Unsweetened Plant-Based Milk Original | 132 | 86 | 114 | 3 | 240 | 317 | 273 | 39 | 33 |
| | | [ab]Sproud - Plant-Based Milk Original | 143 | 100 | 143 | 2 | 240 | 344 | 344 | 75 | 75 |
| | | [ab]Not Milk - Plant-Based Milk Alternative 2% Reduced Fat | 157 | 100 | 157 | 2 | 240 | 377 | 377 | 98 | 98 |

| | | | | | | | | | | | |
|---|---|---|---|---|---|---|---|---|---|---|---|
| | Animal | Kemps - 2% Reduced Fat Milk | 75 | 85 | 63 | 3 | 240 | 179 | 152 | 23 | 19 |
| Yogurt & Cheese | Soy | [a]Silk - Dairy-Free Yogurt Vanilla | 125 | 74 | 93 | 4 | 150 | 188 | 139 | 33 | 24 |
| | | Field Roast - Vegan Chao Shreds Creamy Original | 5 | 0 | 0 | 0 | 28 | 1 | 0 | 0 | 0 |
| | | Field Roast - Vegan Chao Slices Creamy Original | 7 | 0 | 0 | 0 | 20 | 1 | 0 | 0 | 0 |
| | Pulse | Miyoko's - Cultured Vegan Cheese Farmhouse Cheddar | 141 | 64 | 90 | 9 | 28 | 39 | 25 | 15 | 10 |
| | | Daiya - Cheddar Style Slices | 345 | 62 | 214 | 1 | 22 | 76 | 47 | 314 | 195 |
| | Animal | Yoplait - Original French Vanilla | 105 | 81 | 86 | 3 | 170 | 179 | 145 | 33 | 27 |
| | | Kraft - Sharp Cheddar Shredded Cheese | 402 | 89 | 359 | 21 | 28 | 113 | 100 | 19 | 17 |
| | | [a]Kraft Singles - American | 1223 | 93 | 1142 | 17 | 19 | 232 | 217 | 73 | 68 |
| Sausage & Bacon | Soy | Gardein - Plant-Based Sliced Italian Saus'ge | 207 | 59 | 123 | 19 | 56 | 116 | 69 | 11 | 6 |

| | | | | | | | | | | | |
|---|---|---|---|---|---|---|---|---|---|---|---|
| | | MorningStar Farms - Veggie Sausage Links | 80 | 96 | 77 | 18 | 45 | 36 | 34 | 4 | 4 |
| | | [ab]MorningStar Farms - Veggie Bacon Strips | 293 | 100 | 293 | 21 | 16 | 47 | 47 | 14 | 14 |
| | Pulse | Beyond Meat - Beyond Sausage Plant-Based Links Brat Original | 264 | 47 | 124 | 24 | 76 | 201 | 94 | 11 | 5 |
| | | [a]Raised & Rooted - Bratwurst Plant Based Sausage | 196 | 60 | 118 | 20 | 98 | 192 | 116 | 10 | 6 |
| | | Beyond Meat - Beyond Sausage Plant-Based Links Hot Italian | 186 | 71 | 132 | 15 | 76 | 141 | 100 | 12 | 9 |
| | | Beyond Meat - Beyond Breakfast Sausage Plant-Based Links Classic | 267 | 46 | 123 | 21 | 46 | 123 | 56 | 13 | 6 |
| | | Lightlife - Plant-Based Breakfast Patties | 186 | 62 | 116 | 20 | 59 | 110 | 68 | 10 | 6 |
| | Animal | [b]Johnsonville - Original Bratwurst | 122 | 100 | 122 | 18 | 82 | 100 | 100 | 7 | 7 |

| | | | | | | | | | | | |
|---|---|---|---|---|---|---|---|---|---|---|---|
| | | Johnsonville - Italian Sausage Mild | 143 | 93 | 133 | 22 | 56 | 80 | 75 | 7 | 6 |
| | | [ab]Jimmy Dean - Original Pork Sausage Patties | 173 | 100 | 173 | 16 | 68 | 118 | 118 | 11 | 11 |
| | | [b]Hormel - Black Label Original Bacon | 237 | 100 | 237 | 33 | 19 | 45 | 45 | 7 | 7 |
| Chicken & Turkey | Soy | Wholesome Provisions - Just Like Chicken Textured Vegetable Protein | 196 | 49 | 97 | 14 | 49 | 96 | 47 | 14 | 7 |
| | | [a]MorningStar Farms - Original Chik Patties | 184 | 86 | 158 | 15 | 71 | 131 | 112 | 13 | 11 |
| | | [b]Tofurky - Plant Based Deli Slices Oven Roasted | 146 | 100 | 146 | 28 | 52 | 76 | 76 | 5 | 5 |
| | Pulse | [a]Raised & Rooted - Plant Based Nuggets | 156 | 76 | 118 | 8 | 90 | 140 | 106 | 20 | 15 |
| | | [a]Beyond Meat - Beyond Chicken Plant Based Breaded Tenders | 116 | 80 | 93 | 15 | 80 | 93 | 75 | 8 | 6 |
| | Animal | [b]Tyson - Boneless | 260 | 100 | 260 | 17 | 112 | 291 | 291 | 15 | 15 |

| | | | | | | | | | | | |
|---|---|---|---|---|---|---|---|---|---|---|---|
| | | Skinless Chicken Breasts | | | | | | | | | |
| | | [b]Tyson - Chicken Patties | 125 | 100 | 125 | 13 | 76 | 95 | 95 | 10 | 10 |
| | | [a]Hillshire Farm – Oven Roasted Turkey Breast | 273 | 99 | 270 | 15 | 56 | 153 | 151 | 18 | 18 |
| Natural Forms & Traditional Processed Plant-Based Protein Products | Soy | Tofu | 145 | 48 | 69 | 14 | 84 | 122 | 58 | 10 | 5 |
| | | LightLife Tempeh | 260 | 58 | 151 | 24 | 84 | 219 | 127 | 11 | 6 |
| | | Bird's Eye Edamame | 172 | 63 | 109 | 13 | 74 | 128 | 81 | 13 | 8 |
| | | Seapoint Farms Edamame Dry Roasted | 571 | 61 | 348 | 46 | 45 | 257 | 156 | 12 | 8 |
| | | Soymerica Soybeans | 186 | 51 | 94 | 15 | 86 | 160 | 81 | 12 | 6 |
| | Pulses | [b]Chickpeas (canned) | 52 | 100 | 52 | 6 | 122 | 64 | 64 | 8 | 8 |
| | | Chickpeas (Dry) | 104 | 68 | 71 | 8 | 82 | 85 | 58 | 14 | 9 |
| | | Green Lentils (Dry) | 128 | 91 | 116 | 7 | 99 | 127 | 115 | 18 | 16 |
| | | Yellow Split Peas (Dry) | 130 | 76 | 98 | 11 | 98 | 127 | 96 | 12 | 9 |
| | | [b]Fava Bean (Canned) | 73 | 100 | 73 | 8 | 132 | 97 | 97 | 9 | 9 |
| | | Fava Bean (Dry) | 166 | 61 | 101 | 13 | 85 | 141 | 86 | 13 | 8 |
| | | Mung Bean (Dry) | 121 | 78 | 95 | 11 | 101 | 123 | 96 | 11 | 9 |
| | Peanut | Jif Peanut Butter (creamy) | 324 | 70 | 226 | 27 | 33 | 107 | 75 | 12 | 8 |

| | | | | | | | | | | |
|---|---|---|---|---|---|---|---|---|---|---|
| | Soy | WowButter Soy Spread (creamy) | 349 | 88 | 309 | 24 | 32 | 112 | 99 | 15 | 13 |
| | Wheat | Barilla Pasta | 65 | 95 | 62 | 7 | 112 | 73 | 70 | 9 | 9 |
| | Soy | Simply Nature Soybean Spaghetti | 204 | 47 | 97 | 16 | 112 | 229 | 108 | 13 | 6 |
| | Pulse | Banza Spaghetti | 100 | 77 | 77 | 7 | 112 | 112 | 86 | 14 | 11 |

**Supplementary Table 3. Bioaccessible phosphorus and bioaccessible phosphorus-to-protein ratio.** Average total and bioaccessible phosphorus are shown in mg/100g and in mg/serving along with percent bioaccessible phosphorus for each product evaluated. Average total protein in g/100g and total phosphorus-to-protein ratio and bioaccessible phosphorus-to-protein ratio are also shown.

[a] Contains at least one inorganic phosphate additive as identified on the ingredient list.

[b] Percent bioaccessible phosphorus set to the highest possible value of 100% due to calculated value being greater than 100%. Products and the calculated value over 100% are shown below:

- Sproud - Plant-Based Milk Original: 118%
- Not Milk - Plant-Based Milk Alternative 2% Reduced Fat: 106%
- MorningStar Farms - Veggie Bacon Strips: 108%
- Johnsonville - Original Bratwurst: 108%
- Jimmy Dean - Original Pork Sausage Patties: 103%
- Hormel - Black Label Original Bacon: 111%
- Tofurky - Plant Based Deli Slices Oven Roasted: 104%
- Tyson - Boneless Skinless Chicken Breasts: 107%
- Tyson - Chicken Patties: 125%
- Chickpeas (canned): 102%
- Fava Bean (Canned): 102%

**Supplementary Table 4. Total mineral (calcium, magnesium, potassium, and sodium)**

| Food Category | Protein Category | Product | Total Ca (mg/100g) | Total Mg (mg/100g) | Total K (mg/100g) | Total Na (mg/100g) |
|---|---|---|---|---|---|---|
| Beef | Soy | Impossible - Plant-Based Burger Ground | 154 | 57 | 443 | 346 |
| | | Incogmeato MorningStar Farms - Burger Patties | 162 | 83 | 523 | 303 |
| | | Gardein - Plant-Based Ground Be'f | 110 | 72 | 464 | 290 |
| | | MorningStar Farms - Vegan Meat Lovers (Patties) | 49 | 28 | 116 | 473 |
| | Pulse | Beyond Meat - Beyond Beef Beefy Crumbles | 84 | 19 | 214 | 256 |
| | | Sweet Earth - Awesome Grounds | 143 | 45 | 295 | 411 |
| | | Beyond Meat - Beyond Beef Plant Based Patties | 93 | 35 | 252 | 369 |
| | | Gardein - Ultimate Plant-Based Burger | 105 | 35 | 78 | 356 |
| | | Dr. Praeger's - All American Plant-Based Burgers | 95 | 37 | 114 | -- |
| | Animal | Bubba Burger - Original Burger | 6 | 15 | 151 | 57 |
| | | BallPark - Beef Patty | 20 | 14 | 173 | 563 |
| Milk | Soy | Silk - Original Soymilk | 161 | 17 | 139 | 36 |
| | | Pacific Foods - Ultra Soy Plant-Based Beverage Original | 109 | 20 | 192 | 55 |
| | | Silk - Ultra Original Soy Protein Beverage | 166 | 16 | 233 | 111 |
| | Pulse | Ripple - Unsweetened Plant-Based Milk Original | 165 | 5 | 177 | 54 |
| | | Sproud - Plant-Based Milk Original | 113 | 1 | 222 | 76 |

| | | | | | | |
|---|---|---|---|---|---|---|
| | | Not Milk - Plant-Based Milk Alternative 2% Reduced Fat | 56 | 5 | 191 | 75 |
| | Animal | Kemps - 2% Reduced Fat Milk | 84 | 7 | 126 | 33 |
| Yogurt & Cheese | Soy | Silk - Dairy-Free Yogurt Vanilla | 121 | 19 | 175 | 66 |
| | | Field Roast - Vegan Chao Shreds Creamy Original | 6 | 1 | 29 | 832 |
| | | Field Roast - Vegan Chao Slices Creamy Original | 8 | 1 | 22 | 752 |
| | Pulse | Miyoko's - Cultured Vegan Cheese Farmhouse Cheddar | 393 | 38 | 314 | 871 |
| | | Daiya - Cheddar Style Slices | 647 | 7 | 66 | 840 |
| | Animal | Yoplait - Original French Vanilla | 138 | 9 | 112 | 45 |
| | | Kraft - Sharp Cheddar Shredded Cheese | 528 | 19 | 77 | 604 |
| | | Kraft Singles - American | 1331 | 24 | 280 | 604 |
| Sausage & Bacon | Soy | Gardein - Plant-Based Sliced Italian Saus'ge | 87 | 62 | 386 | 665 |
| | | MorningStar Farms - Veggie Sausage Links | 37 | 21 | 8 | 760 |
| | | MorningStar Farms - Veggie Bacon Strips | 128 | 24 | 222 | 1863 |
| | Pulse | Beyond Meat - Beyond Sausage Plant-Based Links Brat Original | 65 | 30 | 317 | 602 |
| | | Raised & Rooted - Bratwurst Plant Based Sausage | 173 | 12 | 665 | 606 |
| | | Beyond Meat - Beyond Sausage Plant-Based Links Hot Italian | 45 | 21 | 15 | 461 |

| | | | | | | |
|---|---|---|---|---|---|---|
| | | Beyond Meat - Beyond Breakfast Sausage Plant-Based Links Classic | 144 | 40 | 444 | 421 |
| | | Lightlife - Plant-Based Breakfast Patties | 53 | 19 | 14 | 409 |
| | Animal | Johnsonville - Original Bratwurst | 9 | 14 | 223 | 691 |
| | | Johnsonville - Italian Sausage Mild | 14 | 19 | 277 | 716 |
| | | Jimmy Dean - Original Pork Sausage Patties | 22 | 14 | 606 | 602 |
| | | Hormel - Black Label Original Bacon | 13 | 22 | 475 | 1483 |
| Chicken & Turkey | Soy | Wholesome Provisions - Just Like Chicken Textured Vegetable Protein | 92 | 89 | 571 | 30 |
| | | MorningStar Farms - Original Chik Patties | 61 | 43 | 427 | 377 |
| | | Tofurky - Plant Based Deli Slices Oven Roasted | 89 | 35 | 655 | 667 |
| | Pulse | Raised & Rooted - Plant Based Nuggets | 139 | 14 | 137 | 640 |
| | | Beyond Meat - Beyond Chicken Plant Based Breaded Tenders | 29 | 24 | 272 | 583 |
| | Animal | Tyson - Boneless Skinless Chicken Breasts | 18 | 31 | 338 | 120 |
| | | Tyson - Chicken Patties | 18 | 17 | 202 | 437 |
| | | Hillshire Farm – Oven Roasted Turkey Breast | 18 | 12 | 462 | 1034 |
| Natural Forms & Traditiona | Soy | Tofu | 111 | 87 | 109 | 20 |
| | | LightLife Tempeh | 97 | 87 | 410 | 20 |
| | | Bird's Eye Edamame | 73 | 83 | 527 | 31 |
| | | Seapoint Farms Edamame Dry Roasted | 138 | 194 | 1371 | 348 |
| | | Soymerica Soybeans | 76 | 73 | 302 | 12 |
| | Pulses | Chickpeas (canned) | 34 | 19 | 139 | 207 |

| | | | | | | |
|---|---|---|---|---|---|---|
| 1 | | Chickpeas (Dry) | 35 | 47 | 169 | 10 |
| Processed Plant-Based Protein Products | | Green Lentils (Dry) | 27 | 30 | 182 | 4 |
| | | Yellow Split Peas (Dry) | 23 | 43 | 271 | 11 |
| | | Fava Bean (Canned) | 46 | 23 | 187 | 496 |
| | | Fava Bean (Dry) | 30 | 43 | 298 | 23 |
| | | Mung Bean (Dry) | 32 | 57 | 273 | 14 |
| | Peanut | Jif Peanut Butter (creamy) | 54 | 157 | 525 | 392 |
| | Soy | WowButter Soy Spread (creamy) | 160 | 138 | 929 | 300 |
| | Wheat | Barilla Pasta | 15 | 28 | 41 | 5 |
| | Soy | Simply Nature Soybean Spaghetti | 83 | 65 | 378 | 23 |
| | Pulse | Banza Spaghetti | 25 | 34 | 144 | 18 |

**Supplementary Table 4. Total mineral (calcium, magnesium, potassium, and sodium).** Average total calcium, magnesium, potassium, and sodium in mg/100g for each food product evaluated. Total calcium, magnesium, potassium and sodium were determined from the freeze-dried food samples by MP-AES using detection wavelengths of 393.366 and 616.217 nm (Ca), 766.491 and 769.897 nm (K), 280.271 nm (Mg), and 588.995 and 589.592 nm (Na). For elements with more than one wavelength, the wavelength with the values within the limit of detection were chosen.
